## Supplementary Material for "Bacterial warfare is associated with virulence and antimicrobial resistance"

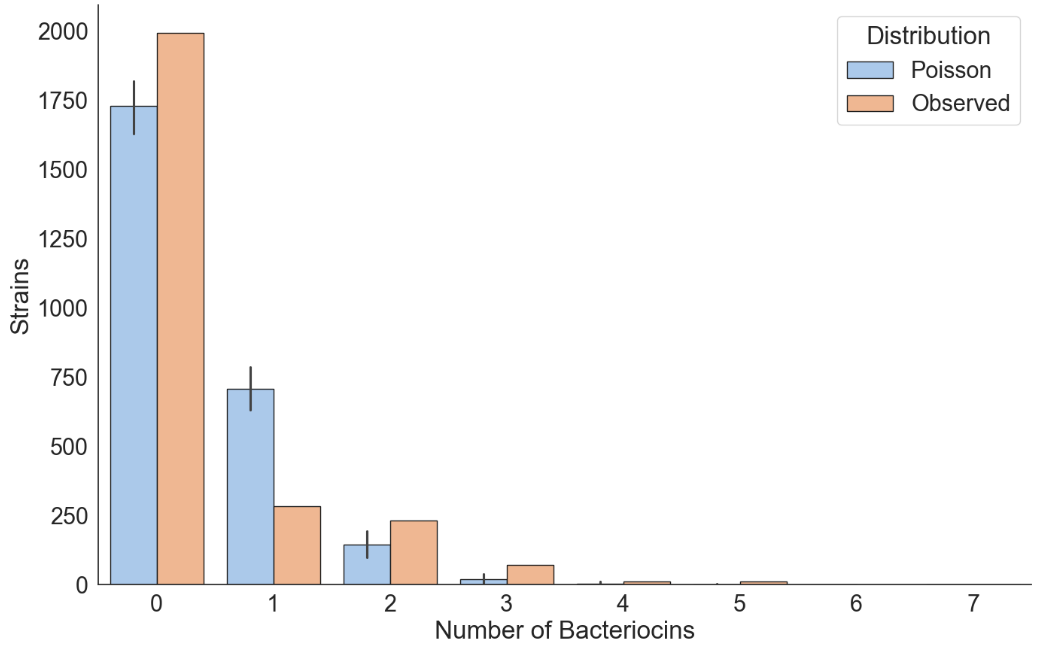


**Supplementary Figure 1. Strains encoding multiple bacteriocins are observed more frequently than predicted by chance.** The observed and theoretical Poisson distribution of bacteriocin distribution amongst strains in the dataset of 2601 *E. coli* genomes. Poisson distribution was calculated using the observed mean number of bacteriocins as λ. The Poisson distribution was approximated for a sample size of 2601 with error bars showing the min and max of 10,000 replicates.

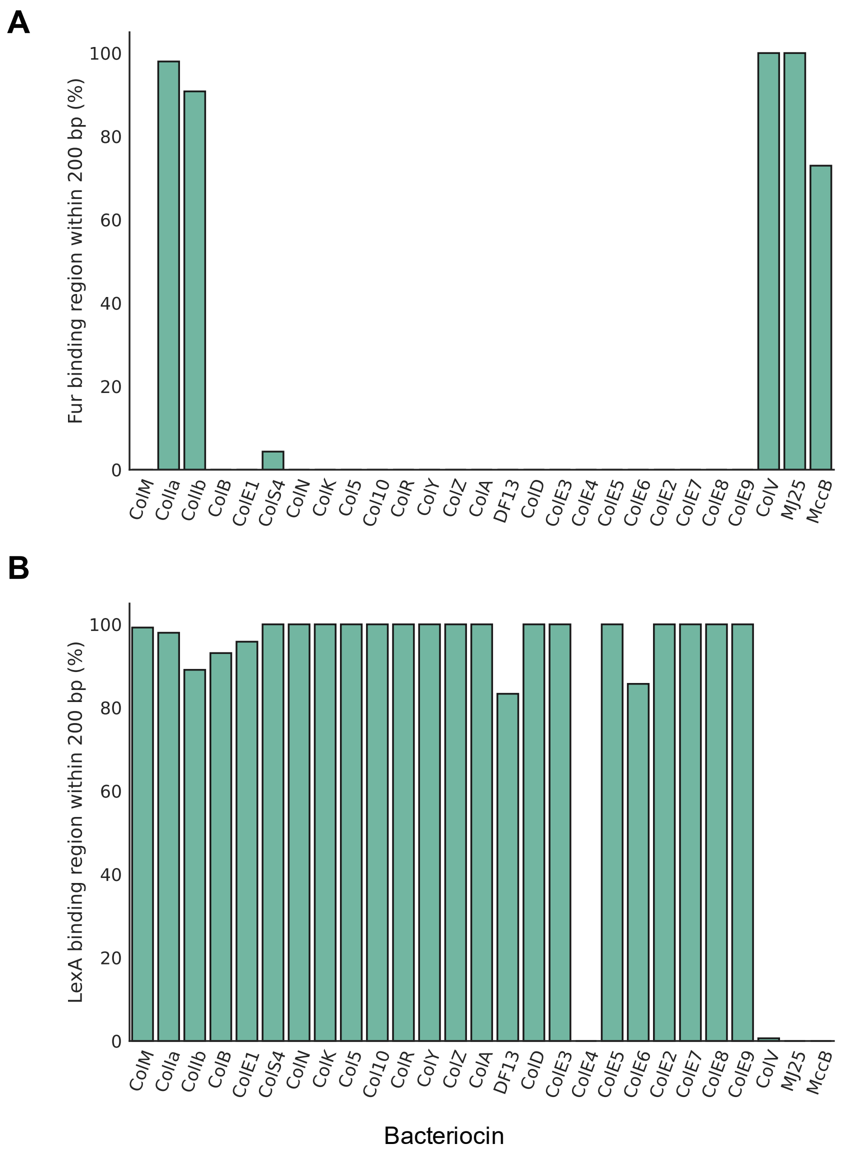


**Supplementary Figure 2. Bacteriocins are regulated by combinations of LexA and Fur.** A) Percentage of bacteriocin genes with an identifiable Fur binding region within 200bp of the predicted start site. B) Percentage of bacteriocin genes with an identifiable LexA binding region within 200bp of the predicted start site.

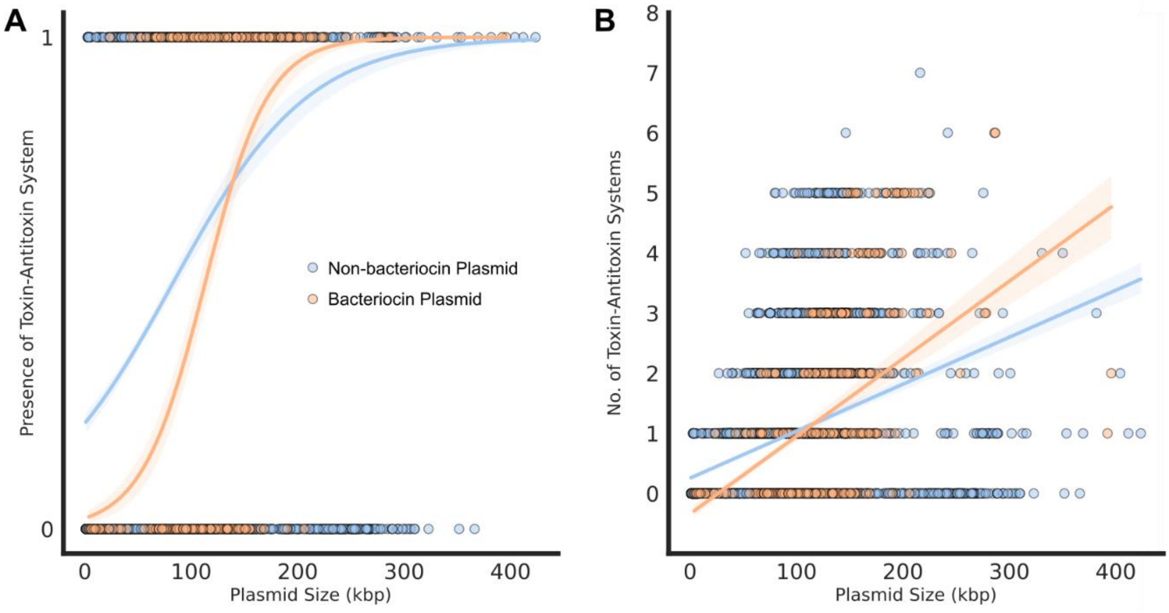


**Supplementary Figure 3. Bacteriocins plasmids are not predicted to be enriched in antimicrobial resistance (AMR) genes compared to the *E. coli* plasmidome when corrected for their large size.** A) Logistic regression model of AMR gene presence/absence of plasmids across the *E. coli* plasmidome. B) Linear regression model of the number of antibiotic class resistances provided by AMR genes in each plasmid as predicted by ResFinder.

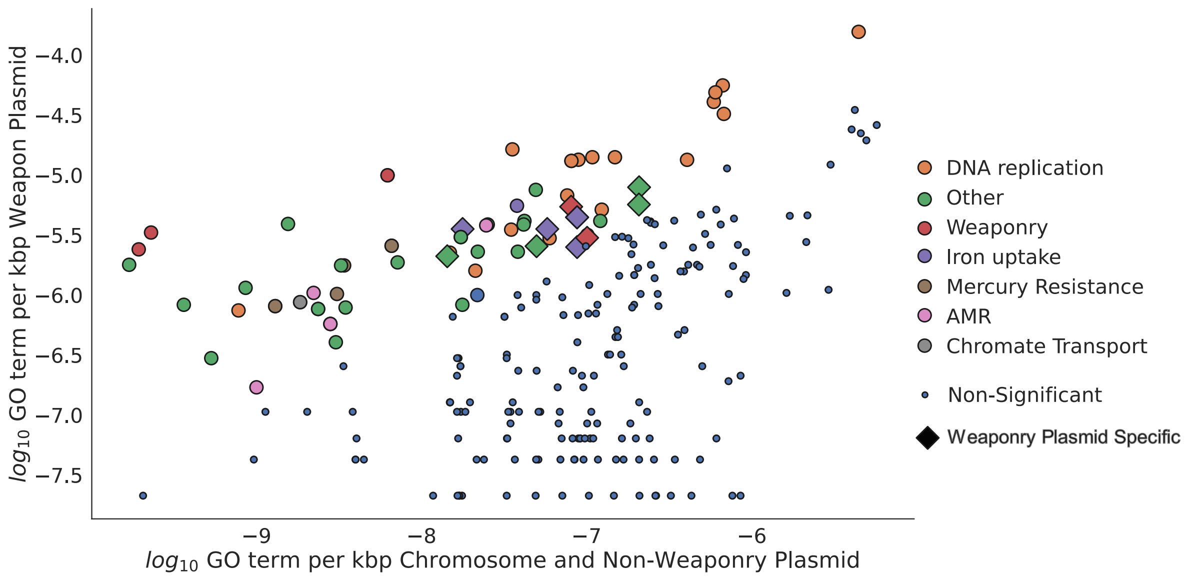


**Supplementary Figure 4. Bacteriocin plasmids bring functions involved in bacterial competition, iron uptake and AMR into a cell.** Analysis of the GO functions of genes in *E. coli*, both on plasmids and chromosomes, demonstrates that bacteriocin plasmids encode functions involved in iron-uptake, competition and AMR.

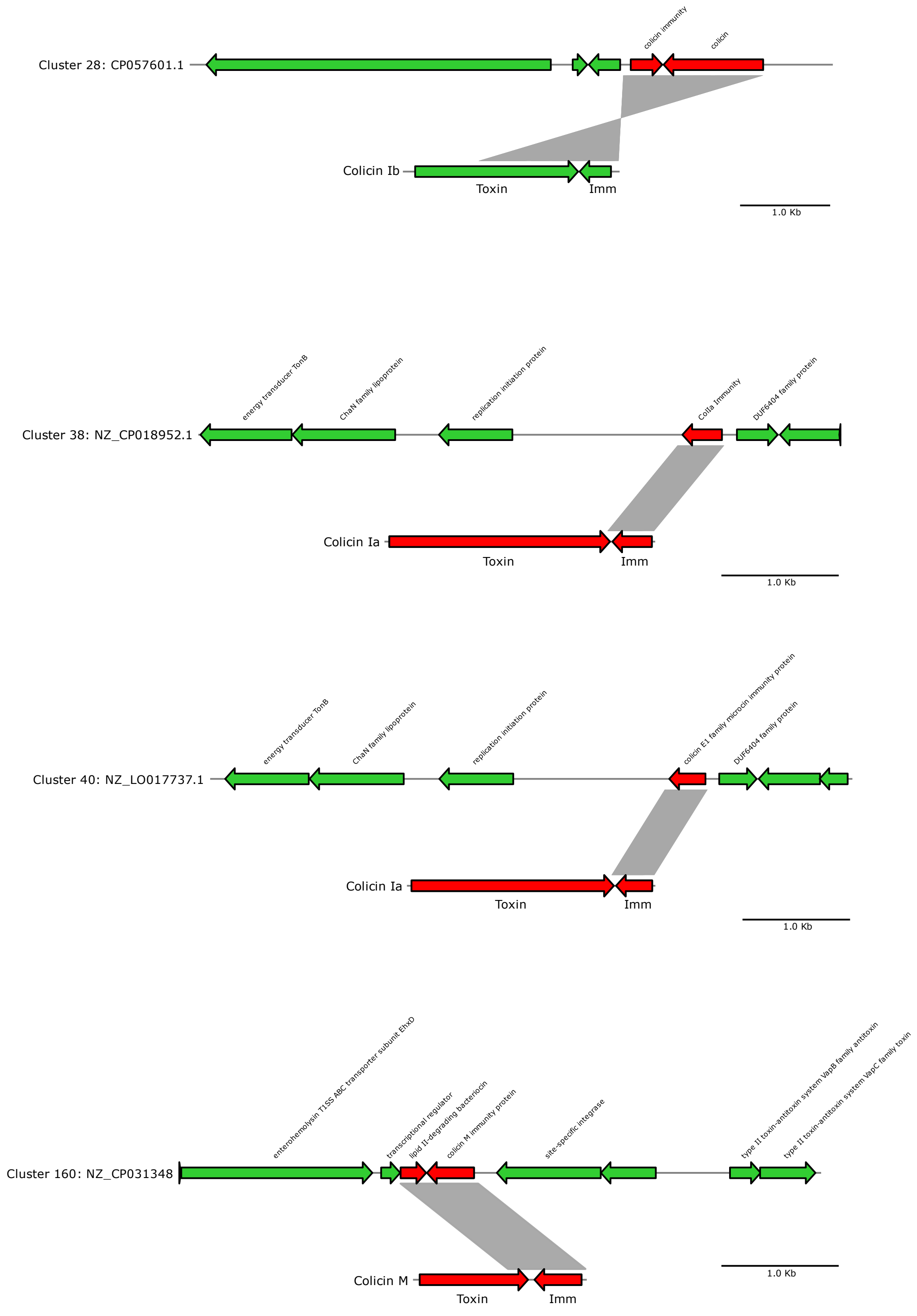


**Supplementary Figure 5. Orphan immunity genes are encoded on multiple plasmids with either no colicin toxin gene or heavily degraded colicin toxin genes.** Alignments of plasmids containing orphan immunity genes with representative colicin-immunity genes. Grey regions indicate alignments as identified by BLASTn with a threshold of 10% identity and 1000bp length.

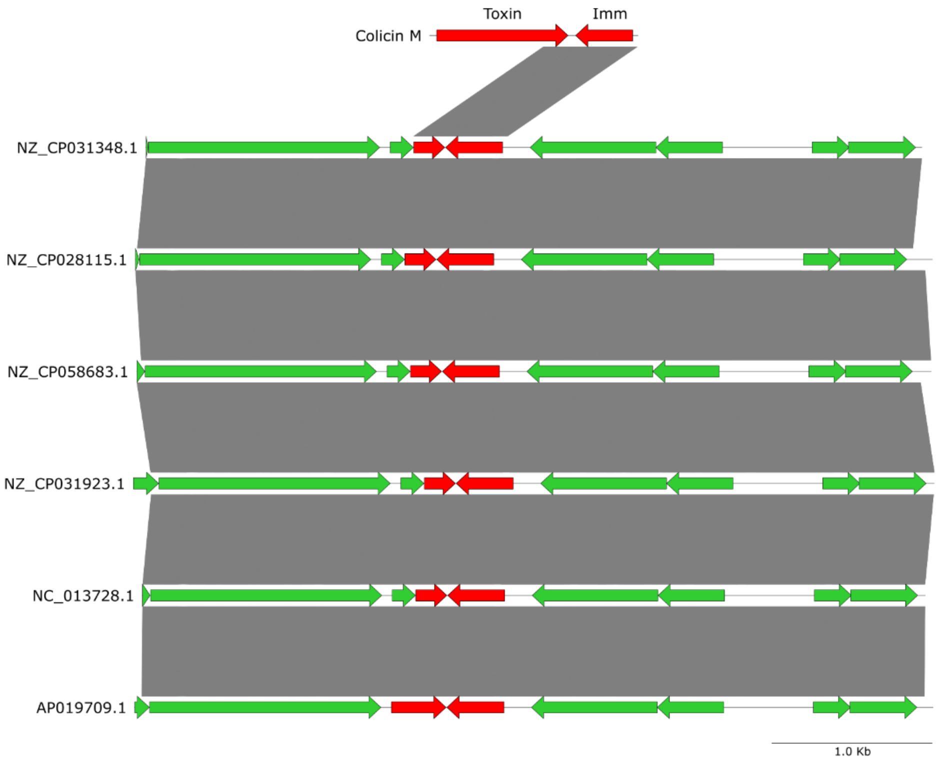


**Supplementary Figure 6. Identical colicin orphan immunity genes and truncated colicins are identified in multiple distinct plasmids within a cluster.** Alignment of the orphan immunity containing regions across multiple plasmids within the same cluster highlights that the immunity gene and truncated colicin are maintained. Plasmids are of different sizes and isolated from different environments (NZ_CP031348.1: Size - 89518 bp, Country of origin – USA, Niche – Cattle feces; NZ_CP028115.1: Size – 90123 bp, Country of origin – USA, Niche – Creek; NZ_CP058683.1: Size – 92578 bp, Country of origin - Germany**,** Niche – Salad; NZ_CP031923.1: Size – 95298 bp, Country of origin – Canada, Niche – Human feces; NC_013728.1: Size – 111481 bp, Country of origin – Australia, Niche – unknown; AP019709.1: Size 86874 bp, Country and origin – Japan, Niche – Human feces.)

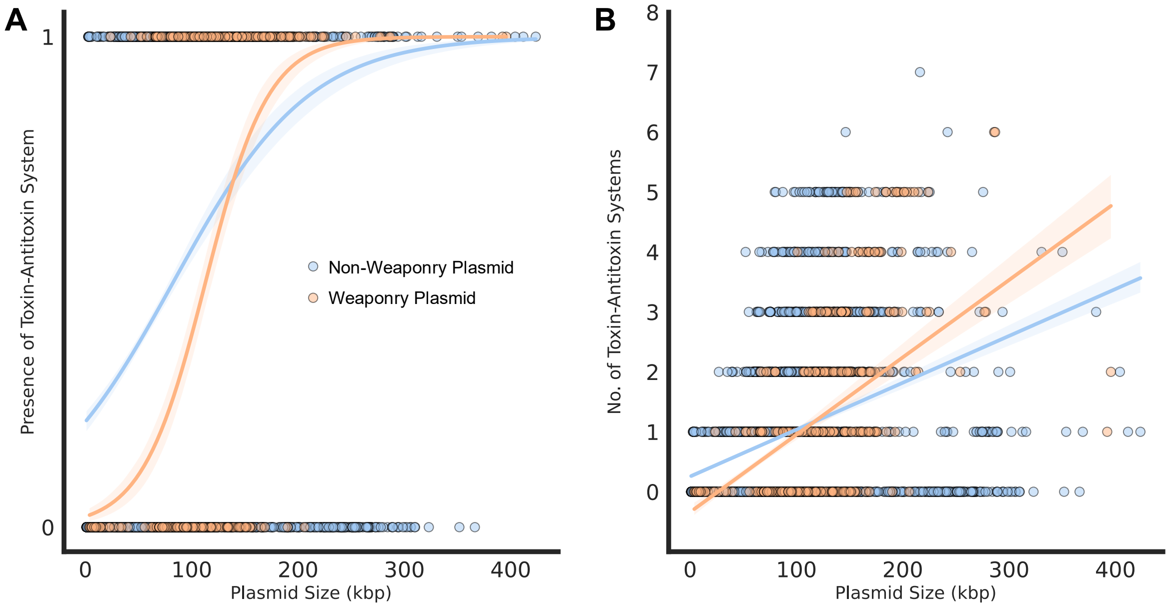


**Supplementary Figure 7. Bacteriocin plasmids are not less likely to encode a toxin-antitoxin system or encode fewer toxin-antitoxin systems.** A) Logistic regression model of the presence of toxin-antitoxin systems on plasmids in the *E. coli* plasmidome. B) Linear regression showing the number of predicted toxin-antitoxin systems on plasmids within the *E. coli* plasmidome accounting for plasmid size.

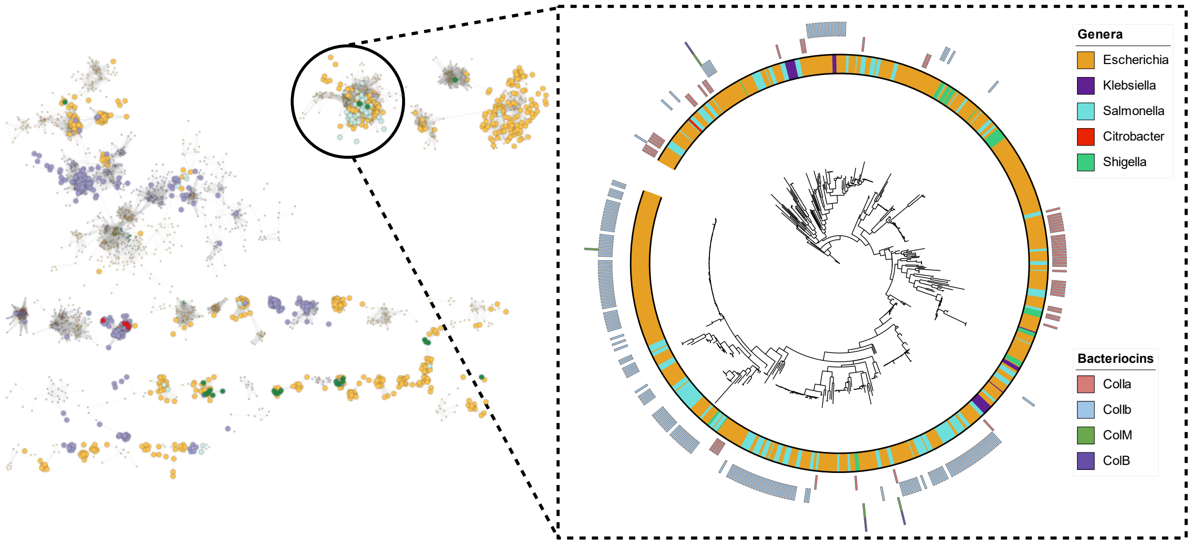


**Supplementary Figure 8. Plasmid clustering reveals similar plasmids shared between *Salmonella*, *E. coli* and *K. pneumoniae*, suggesting that plasmid host range does not limit the sharing of bacteriocin plasmids between the three species.** Analysis of a plasmid cluster which contains bacteriocin and non-bacteriocins plasmids shared between three species shows closely related plasmids are shared between species. Phylogeny was calculated from conserved genes on plasmid backbones.

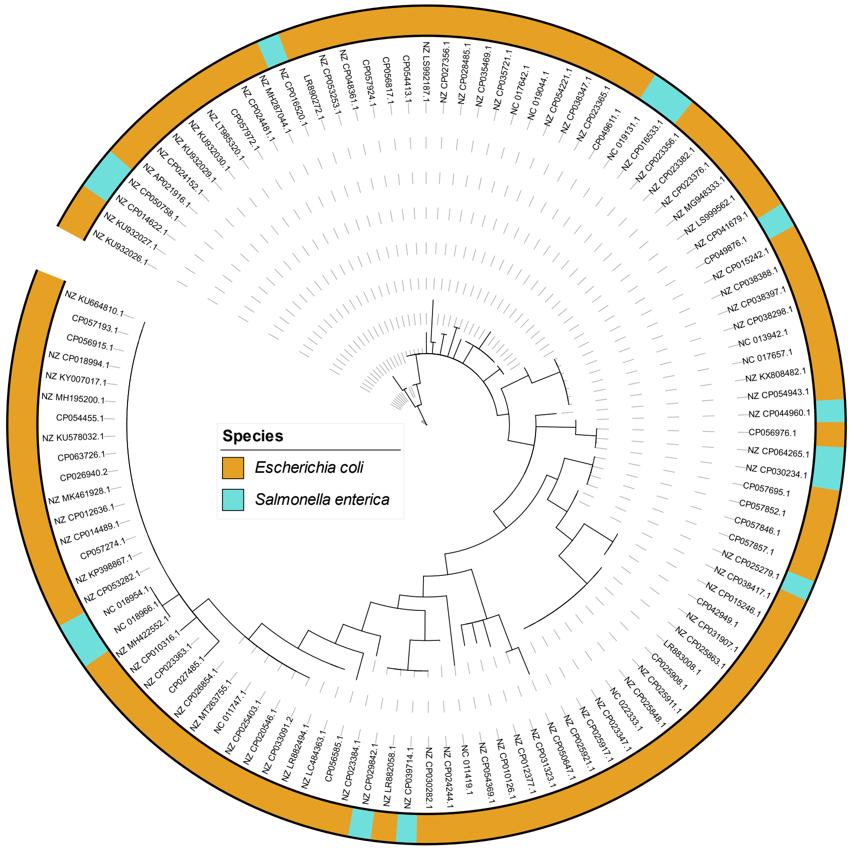


**Supplementary Figure 9. Phylogeny of Colicin Ia suggests genes have been shared multiple times between *Salmonella* and *E. coli*.** Phylogeny of concatenated Colicin Ia and immunity genes from *E. coli* and *Salmonella*.

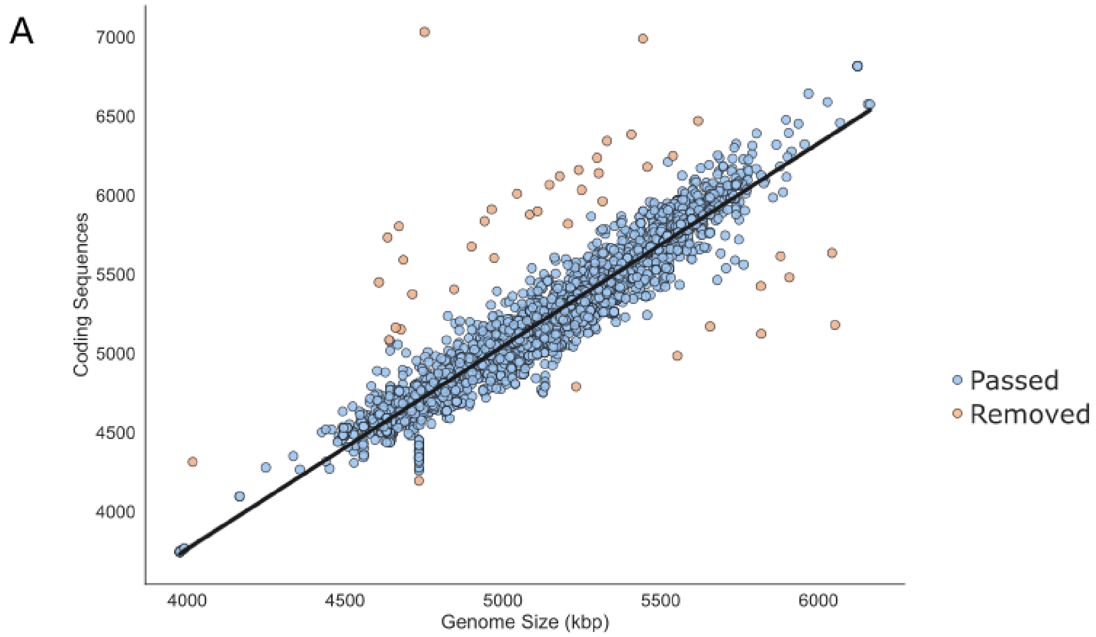


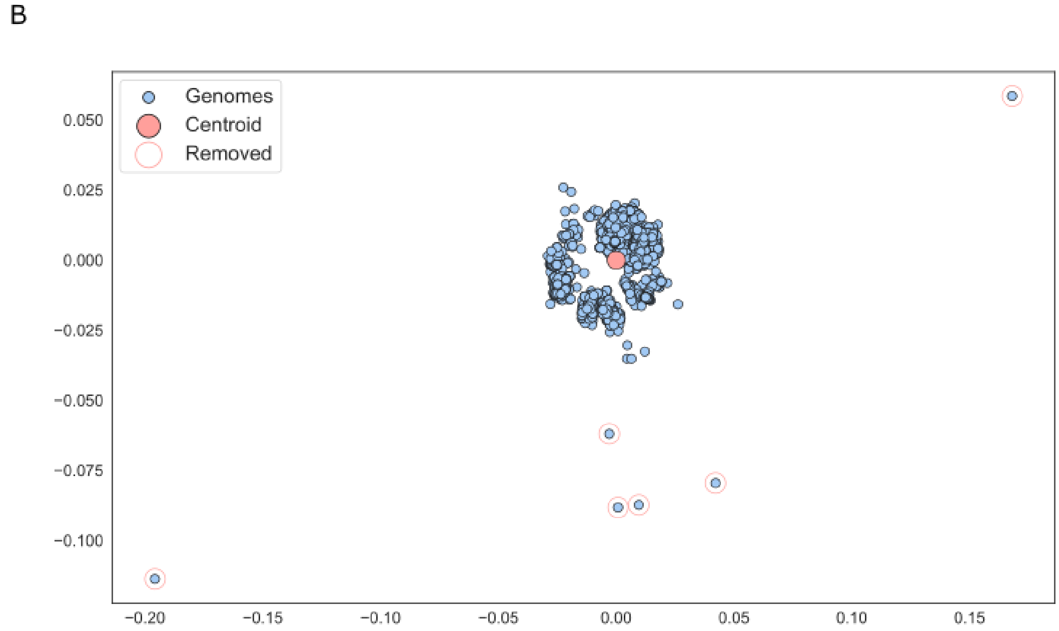


**Supplementary Figure 10. Pre-processing of *E. coli* pangenome strains to remove contamination.** A) *E. coli* genomes which deviated from a linear relationship between genome length and number of coding sequences were removed from further analysis. B) Possible contaminated genomes were removed by analysis of MASH distances between all strains in the population. After MDS scaling, outlier strains were removed.

**Supplementary Table 1. Sequences used to identify bacteriocins in *E. coli*.**

| Bacteriocin Gene | Accession |
|---|---|
| ColM | WP_000449473.1 |
| ColM immunity | WP_105457450.1 |
| ColIa | WP_001582575.1 |
| ColIa Immunity | WP_001080729.1 |
| ColIb | WP_010891256.1 |
| ColIb Immunity | WP_000762570.1 |
| ColB | WP_021539362.1 |
| ColB Immunity | WP_032084022.1 |
| ColE1 | P02978.1 |
| ColE1 Immunity | WP_000058760.1 |
| ColS4 | CAB46008.1 |
| ColS4 Immunity | WP_104807860.1 |
| ColK | WP_159420795.1 |
| ColK Immunity | WP_011264161.1 |
| Col5 | WP_000362089.1 |
| Col5 Immunity | WP_011264161.1 |
| Col10 | Q47125.1 |
| Col10 Immunity | MCD4231352.1 |
| ColR | WP_104355620.1 |
| ColR Immunity | WP_040090309.1 |
| ColY | WP_010892543.1 |
| ColY Immunity | WP_021534853.1 |
| ColZ | WP_172694321.1 |
| ColA | WP_008323639.1 |
| ColA Immunity | EEW1531982.1 |
| CloDF13 | WP_010891190.1 |
| CloDF13 Immunity | RRE37366.1 |
| ColD | WP_016245160.1 |
| ColD Immunity | WP_001038407.1 |
| ColE3 | WP_000012964.1 |
| ColE3 Immunity | WP_000523346.1 |
| ColE4 | CAA45167.1 |
| ColE4 Immunity | RRE37366.1 |
| ColE5 | WP_015420185.1 |
| ColE5 Immunity | ELC99665.1 |
| ColE6 | WP_052980126.1 |
| ColE6 Immunity | WP_001419702.1 |
| ColE2 | WP_172690082.1 |
| ColE2 Immunity | WP_000421100.1 |
| ColE7 | WP_021530049.1 |
| ColE7 Immunity | WP_000420692.1 |
| ColE8 | WP_012766032.1 |
| ColE8 Immunity | WP_000421100.1 |
| ColE9 | WP_012644886.1 |

| | |
|---|---|
| ColE9 Immunity | WP_012644887.1 |
| mchA | AJ009631 |
| mchS1 | AJ009631 |
| mchS2 | AJ009631 |
| mchS3 | AJ009631 |
| mchS4 | AJ009631 |
| mchX | AJ009631 |
| mchI | AJ009631 |
| mchB | AJ009631 |
| mchC | AJ009631 |
| mchD | AJ009631 |
| mchE | AJ009631 |
| mchF | AJ009631 |
| mchI | AJ009631 |
| mcmI | AJ515252 |
| mcmA | AJ515253 |
| mcmM | AJ515254 |
| mcbA | FM877811 |
| mcbB | FM877811 |
| mcbC | FM877811 |
| mcbD | FM877811 |
| mcbE | FM877811 |
| mcbF | FM877811 |
| mcbG | FM877811 |
| mcjA | AF061787 |
| mcjB | AF061787 |
| mcjC | AF061787 |
| mcjD | AF061787 |
| cvaC | AF062848 |
| cvaI | CP005931 |
| cvaA | CP005931 |
| cvAB | CP005931 |

**Supplementary Table 2. Additional sequences used to identify Colicin-like proteins in *Escherichia, Klebsiella, Salmonella, Citrobacter* and *Shigella*.**

| Colicin-Like protein | Genus/Species of origin | Accession |
|---|---|---|
| SalE2 | *Salmonella* | KTM78572.1 |
| SalE3 | *Salmonella* | GAS18013.1 |
| SalE7 | *Salmonella* | KSU39545.1 |
| SalIa | *Salmonella* | OIN35410.1 |
| SalIb | *Salmonella* | OIN32443.1 |
| ColA-like protein_KP | *K. pneumoniae* | SAV78255.1 |
| Colicin-like protein | *K. aerogenes* | WP_063414841.1 |
| Colicin-like protein | *K. oxytoca* | WP_024273778.1 |
| Colicin-like protein | *K. variicola* | KDL88409.1 |
| Colicin-like protein | *K. pneumoniae* | BAS34675.1 |
| Colicin-like protein | *K. pneumoniae* | EWD35590.1 |
| ColM-like protein | *Klebsiella* | WP_047066220.1 |
| ColM-like protein | *K. variicola* | CTQ17225.1 |
| ColM-like-protein | *K. aerogenes* | WP_015367360.1 |
| Klebicin C | *K. penumoniae* | AAT85004.1 |
| Klebicin D | *K. oxytoca* | WP_224261359.1 |
